# T-PGI: an engineered STITCHR system for scarless, programmable genome integration

**DOI:** 10.1101/2025.09.28.679074

**Authors:** Kaveh Daneshvar, Yun Ni, Dev Paudel, Sanah Langer, Hyaeyeong Kim, Sara Ashrafi Kakhki, Omar O. Abudayyeh, Jonathan S. Gootenberg, Jonathan D. Finn, Sandeep Kumar, Joo-Hyung Lee

**Author notes:** These authors contributed equally (Kaveh Daneshvar, Yun Ni, Joo-Hyung Lee). Corresponding authors: Joo-Hyung Lee; Sandeep Kumar, Jonathan D Finn.

## Abstract

Scarless, programmable insertion of defined DNA remains a central goal for therapeutic genome editing. We introduce T-PGI (Transposon-mediated programmable genomic integration), an engineered implementation of STITCHR that preserves target-primed reverse transcription (TPRT) while substantially improving efficiency, specificity, and modularity. T-PGI uses R2Tocc, a low-background ortholog that we further engineered through defined deletions, rational point mutations, and modular domain insertions to enhance performance. The system employs paired nCas9 nicks flanking homology arms and bicistronic co-expression with dual NLSs, together with optimized RNA donor designs. Using combinatorial guide–template screening, T-PGI achieves robust integration across diverse cargos, including precise cassette insertion following multi-kilobase deletions. Short- and long-read sequencing confirm high-fidelity insertion with minimal local indels or structural variants. Collectively, these advances establish T-PGI as a practical and adaptable platform for accurate, scarless genome integration and provide concise design principles spanning enzyme architecture, donor configuration, and guide pairing for next-generation therapeutic editing.

## Introduction

The advent of CRISPR–Cas9–based genome editing has transformed our ability to manipulate the genome with high precision and has spurred the development of a wide array of technologies for targeted modifications (Jinek et al., 2012). While these advances have led to substantial biological and therapeutic insights, current genome editing approaches still face practical challenges, including dependency on cellular repair pathways, variability in editing efficiency, limited control over cargo size and format, and risks of unintended genomic alterations (Kosicki et al., 2018). These limitations have prompted continued efforts to explore alternative strategies that can achieve programmable, scarless, and efficient genome insertion across diverse contexts.

In this regard, the recently developed **<u>s</u>**ite-specific **<u>t</u>**arget-primed **i**nsertion via **t**argeted **<u>C</u>**RISPR **<u>h</u>**oming of **r**etroelements (STITCHR) platform marked a conceptual advance by enabling scarless DNA insertion through target-primed reverse transcription (TPRT), leveraging a fusion of Cas9 nickase and a retrotransposon-derived reverse transcriptase (Fell et al., 2025). This system demonstrated proof-of-concept insertions using either plasmid or RNA-encoded templates in both dividing and non-dividing cells, without relying on endogenous homologous recombination or generating double-strand breaks. However, the initial implementation of STITCHR was limited by relatively low insertion efficiency, especially when compared to established editing platforms, thereby restricting its immediate utility for therapeutic genome engineering.

In this study, we present a comprehensive engineering-focused advancement of the STITCHR platform, aimed at overcoming these early limitations and enhancing its performance and versatility. We systematically optimized core components, including the Cas9 nickase-retrotransposon fusion configuration, nuclear localization strategies, linker composition, and enzyme expression methods, resulting in significantly improved protein activity and cellular delivery. Parallel efforts in donor template design, such as untranslated region (UTR) trimming, homology arm tuning, and post-integration PAM removal, further boosted insertion efficiency. We also performed sgRNA pair screening, uncovering key guide–template combinations that maximized editing outcomes across multiple loci.

Importantly, the engineered STITCHR platform now supports a broad range of editing applications, including long-range deletions coupled with an insertion, precision rewriting with multiple point mutations, and full-length cDNA knock-in, achieving significantly higher insertion efficiency than the original system. These findings not only establish a robust genome insertion platform but also highlight the critical role of platform engineering in advancing emerging editing technologies toward translational use.

## Result

### T-PGI enables targeted, scarless DNA insertion via a TPRT-based mechanism

Transposon-mediated programmable genomic integration (T-PGI) is composed of three core elements: (1) paired Cas9 nickase (nCas9) RNPs with forward and reverse sgRNAs that introduce single-strand nicks at the left and right homology arms, (2) an engineered retrotransposon-derived reverse transcriptase (R2Tocc), and (3) an RNA donor template containing short hairpin structure for RNP assembly and homology arms flanking the desired cargo. As illustrated in **Fig. 1**, nCas9 generates a site-specific nick, exposing a single-stranded region that facilitates hybridization of the donor template via its homology arms. This annealed structure serves as a primer for the R2 enzyme to initiate target-primed reverse transcription (TPRT), copying the donor sequence directly into the genome. This mechanism enables precise, scarless insertion without the need for double-strand breaks. Although the mechanism of second-strand synthesis remains incompletely understood, we expect that it is mediated by host DNA repair machinery, as the R2 enzyme lacks an RNase H domain required for degradation of the RNA strand during reverse transcription (Eickbush & Eickbush, 2015).

**Fig 1.**
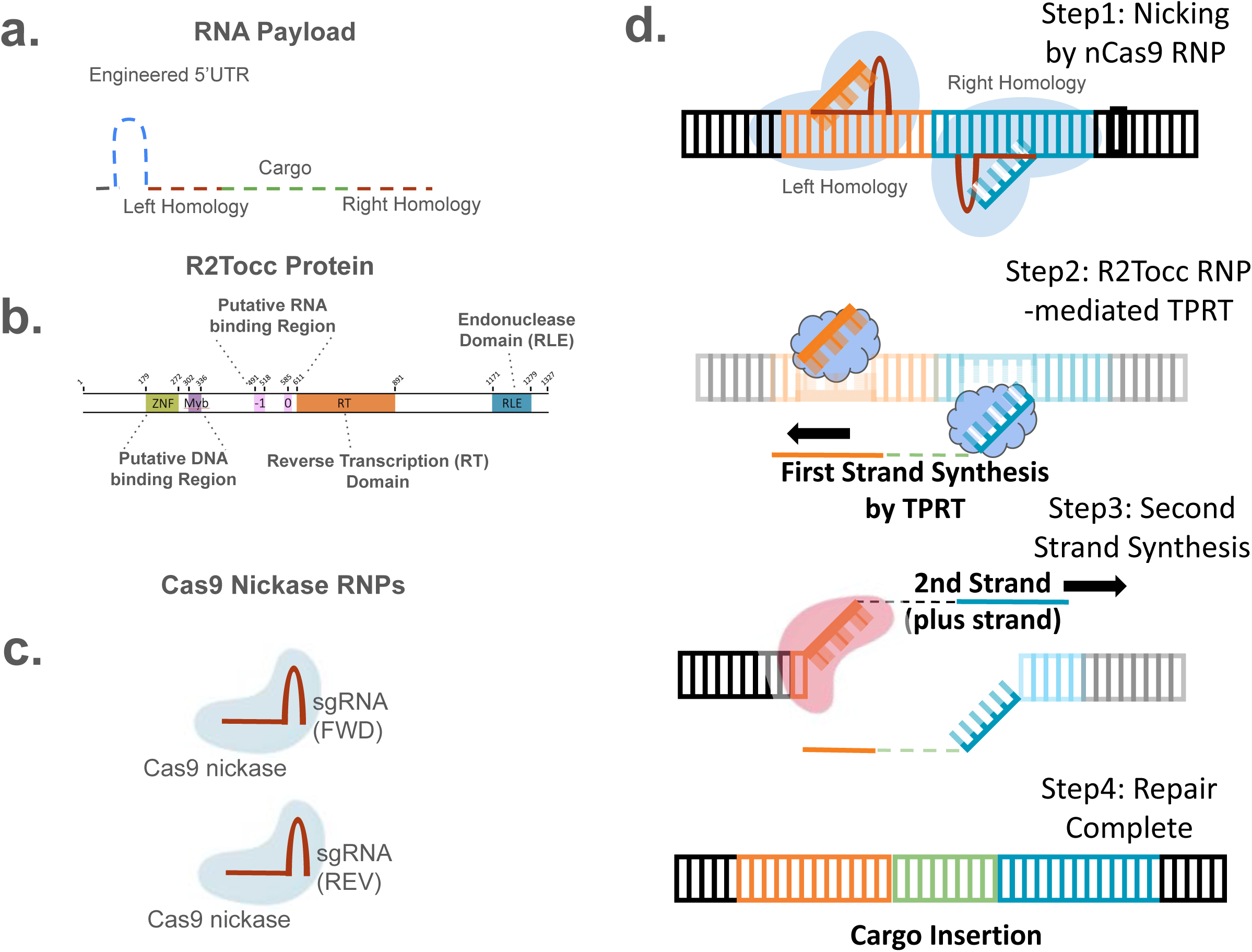
Overview of the T-PGI platform for scarless, programmable DNA insertion. Schematic representation of Transposon-mediated programmable genomic integration (T-PGI)-mediated genome editing. (a) The RNA payload contains an engineered 5′ UTR (blue), target-specific left and right homology sequences (brown), and the cargo sequence (green). (b) The R2Tocc protein includes domains characteristic of R2 retrotransposons: a DNA-binding region (aa 179–336), an RNA-binding region (aa 491–611), a reverse transcriptase domain (aa 611–891), and a restriction-like endonuclease (RLE) domain (aa 1171–1279). (c) Two Cas9 nickase (nCas9) RNP complexes are programmed with forward (FWD) and reverse (REV) sgRNAs to introduce nicks at the left and right homology arms. (d) Mechanism of T-PGI-mediated insertion. Step 1: nCas9 RNPs generate nicks flanking the target site. Step 2: R2Tocc initiates target-primed reverse transcription (TPRT) from the right homology arm to synthesize the first strand. Step 3: The nascent strand anneals to the left nick to initiate second-strand synthesis. Step 4: Repair is completed, resulting in scarless cargo insertion of up to 10.9kb.

Throughout this study, we systematically dissected the contribution of each component, enzyme configuration, donor template structure, and sgRNA design, toward improving STITCHR’s overall performance. The following sections describe how platform-level optimization of these components led to substantial improvements in editing efficiency.

### T-PGI supports broad cargo flexibility and large-scale genomic rewriting

In the original STITCHR study (Fell et al., 2025), R2Tg and R2Tocc emerged as the two most efficient orthologues for TPRT-mediated integration. To directly compare their performance, we conducted side-by-side assays using a shared donor template and a synthetic genomic target, and observed that both enzymes supported similar levels of insertion efficiency (**Supp. Fig.1a**). In the previous findings, R2Tg was shown to retain residual activity at its native 28S rDNA locus, leading to unintended off-target insertions. By contrast, R2Tocc displayed markedly reduced integration at the endogenous 28S site while preserving high reprogramming efficiency (Fell et al., 2025). Given this favorable balance, we selected R2Tocc as the preferred orthologue for subsequent engineering and biotechnological applications.

We next evaluated whether STITCHR could accommodate a wide range of DNA cargo sizes, a key consideration given the mechanistic nature of retrotransposon-mediated integration. Because R2 enzymes operate through reverse transcription, they are known to produce 5’-truncated insertions when the reverse transcriptase prematurely disengages (Eickbush & Eickbush, 2015). Thus, it remained unclear how well TPRT could handle increasingly large DNA templates in mammalian cells, and whether cargo length would inversely impact insertion efficiency. To address this, we evaluated donor templates encoding inserts of 26 bp, 689 bp, and 4.1 kb, spanning a broad range of sequence lengths. Remarkably, STITCHR supported efficient insertion across all cargo sizes with only modest variation (**Supp. Fig.1b**), suggesting that the engineered system effectively stabilizes reverse transcription initiation and elongation even with large payloads. This is consistent with prior observations that retrotransposon-based integration can support high-fidelity insertion of both therapeutic genes and synthetic sequences spanning several kilobases in length (Fell et al., 2025).

To better understand factors limiting short-sequence insertion, we optimized donor template design. When a 38 bp insert was flanked by 50 bp homology arms, insertion efficiency was suboptimal. However, extending the homology arms to 100 bp and including a 200 bp random filler sequence substantially improved performance (**Fig. 2a**). This suggests that very short templates may require additional sequence context to facilitate primer annealing or reverse transcription initiation. Furthermore, eliminating the PAM site post-insertion to prevent re-cleavage by nCas9 further boosted insertion efficiency, highlighting the importance of rational template design (**Fig.2a**).

**Fig 2.**
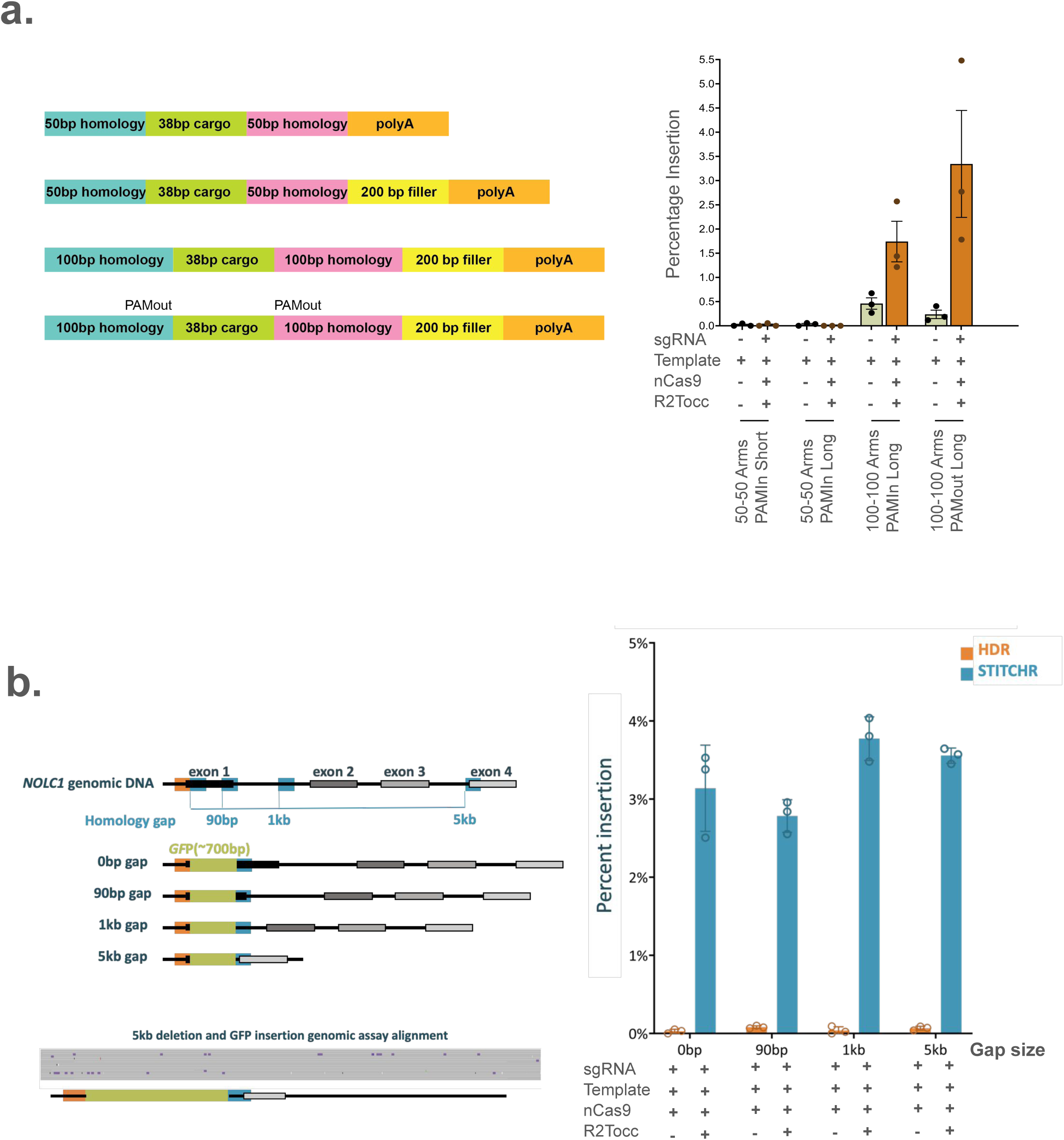
T-PGI supports efficient short cargo insertion and precise rewriting with large deletions. (a) Left: Schematic of donor templates for 38 bp cargo with 50 bp homology arm, adding filler sequences, 100 bp homology arm and PAM sequence removed after integration. Right: ddPCR-based quantification of insertion efficiency at the *NOLC1* locus. Template-only condition was used as a control. (b) T-PGI supports simultaneous long-range deletion and targeted insertion. Left: Schematic of the *NOLC1* locus showing the genomic target sites and predicted insertion outcomes across increasing homology gaps (90 bp, 1 kb, and 5 kb). Right: Quantification of GFP cassette insertion efficiency by ddPCR across deletion lengths. Controls include HDR-like condition. Bottom left: Representative long-read sequencing results (Oxford Nanopore) confirming precise replacement without truncation or aberrant mutations at insertion junctions.

While these results demonstrated that STITCHR is compatible with a broad range of cargo sizes, we next sought to determine whether it could support more complex genome engineering tasks such as large-scale rewriting. The original STITCHR report (Fell et al., 2025) showed efficient replacement of short genomic segments (∼150 bp), but the maximum extent of simultaneous deletion and cassette replacement had not been defined. To address this, we tested whether progressively longer genomic deletions (90 bp, 1 kb, and 5 kb) could be seamlessly replaced with a 689 bp GFP cassette using STITCHR. Remarkably, insertion efficiency remained stable across all deletion contexts, showing no measurable drop compared to non-deletion conditions (**Fig. 2b**). Long-read sequencing further confirmed the sequence integrity of the inserted cassettes, revealing no evidence of partial insertions, unexpected truncations, or aberrant mutations at the junction sites (**Fig. 2b**). These findings indicate that STITCHR can be extended to support long-range editing scenarios such as exon replacement, multi-exon deletion repair, or therapeutic gene rewiring, further expanding its potential utility.

### Insertion fidelity is maintained across diverse editing outcomes

While increasing insertion efficiency is critical, therapeutic genome editing platforms must also ensure high fidelity and minimal genomic disruption. Prior studies of prime editing and other reverse transcriptase–based systems have shown that editing accuracy often declines with increasing cargo size, in part due to premature dissociation of the RT enzyme or low fidelity of the RT process (Baranauskas et al., 2012, Anzalone et al., 2021, Eickbush & Eickbush, 2015). Given that STITCHR also relies on reverse transcription, we assessed the fidelity of long-insert copying during TPRT-mediated integration. Additionally, while STITCHR utilizes Cas9 nickase for targeted DNA cleavage, the R2 enzyme retains intrinsic endonuclease activity.

Consequently, we investigated whether integration events are accompanied by unintended genomic alterations, such as local indels, truncations, or concatemers at or near the insertion site.

To address these questions, we performed orthogonal assessments using both short- and long-read next-generation sequencing. In a precision rewriting experiment, a 90 bp genomic region was replaced using a donor containing nine designed missense mutations. All intended variants were introduced with high accuracy by targeted amplicon sequencing (**Supp. Fig. 2a**). In a broader fidelity assessment, we compared the rewritten 138 bp region to the wild-type reference: 81.4% of reads in the edited population perfectly matched the donor sequence, versus 93.6% in unedited cells. There was no increase in indels or unintended mutations near the homology arms (**Supp. Fig. 2b**).

To extend this analysis to long insertions, we applied PacBio sequencing to STITCHR-edited alleles that incorporated a 689 bp GFP cassette at a 5 kb deletion site. Sequence analysis over a 400 bp window within the inserted region showed 71.9% accuracy, and we found no evidence of structural rearrangements, incomplete insertions, scars, or concatemers (**Supp. Fig. 2c**). Compared to shorter insertions (e.g., 138 bp), the reduced accuracy observed with the 689 bp cassette suggests that insertion fidelity decreases with increasing sequence length. Importantly, this effect does not appear to result from structural variants but is more likely attributable to the intrinsic error rate of reverse transcription. Together, these findings demonstrate that STITCHR enables efficient and accurate sequence insertion across a wide range of edit types and lengths.

### Template design impacts insertion efficiency

We revisited template design principles using the R2Tocc backbone. Inspired by earlier work showing that 5’ and 3’ UTRs influence performance with R2Tg enzyme (Fell et al., 2025), we systematically tested full-length vs. truncated 5’ UTRs and the presence or absence of 3’ UTRs. Consistent with previous observations, truncated 5’ UTRs improved insertion efficiency relative to full-length sequences, while 3’ UTRs were dispensable (**Fig. 3a**). We next examined how varying homology arm (HA) lengths affected efficiency. Templates bearing 50 or 100 bp HAs in various left–right combinations yielded no statistically significant differences in insertion (**Fig. 3b**). However, the use of 100 bp HAs was prioritized in subsequent screens due to increased flexibility in sgRNA positioning, which is particularly advantageous for locus-specific therapeutic designs.

**Fig 3.**
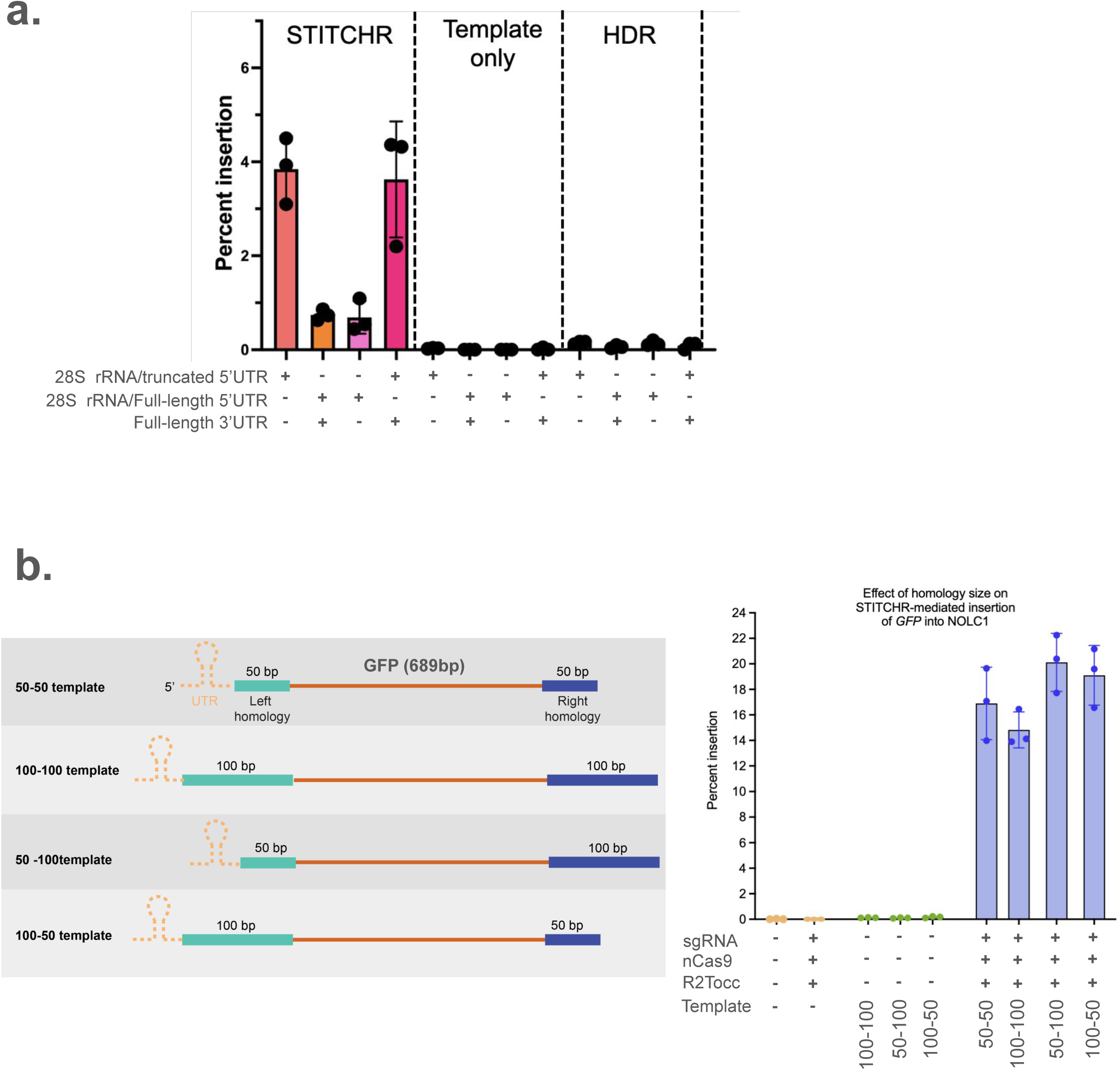
Optimization of UTR and homology arm configurations for T-PGI donor templates. (a) Comparison of insertion efficiency using donor templates with different UTR configurations. Template-only and HDR-like conditions (nCas9 + gRNAs + template) were used as negative controls. (b) Assessment of homology arm (HA) length using templates with 50 bp or 100 bp arms in various combinations. Left: Schematic of donor designs. Right: ddPCR-based quantification of GFP insertion at the *NOLC1* locus for each template configuration.

### Guide RNA pairing and screening to optimize T-PGI targeting

Since T-PGI relies on single-strand breaks generated by nCas9, the precise positioning and combination of sgRNAs flanking the homology arms are likely to impact overall editing efficiency. We first tested single-guide versus dual-guide configurations, targeting either the left or right arm, or both simultaneously. Dual-guide use significantly improved editing performance (**Fig. 4a**).

**Fig 4.**
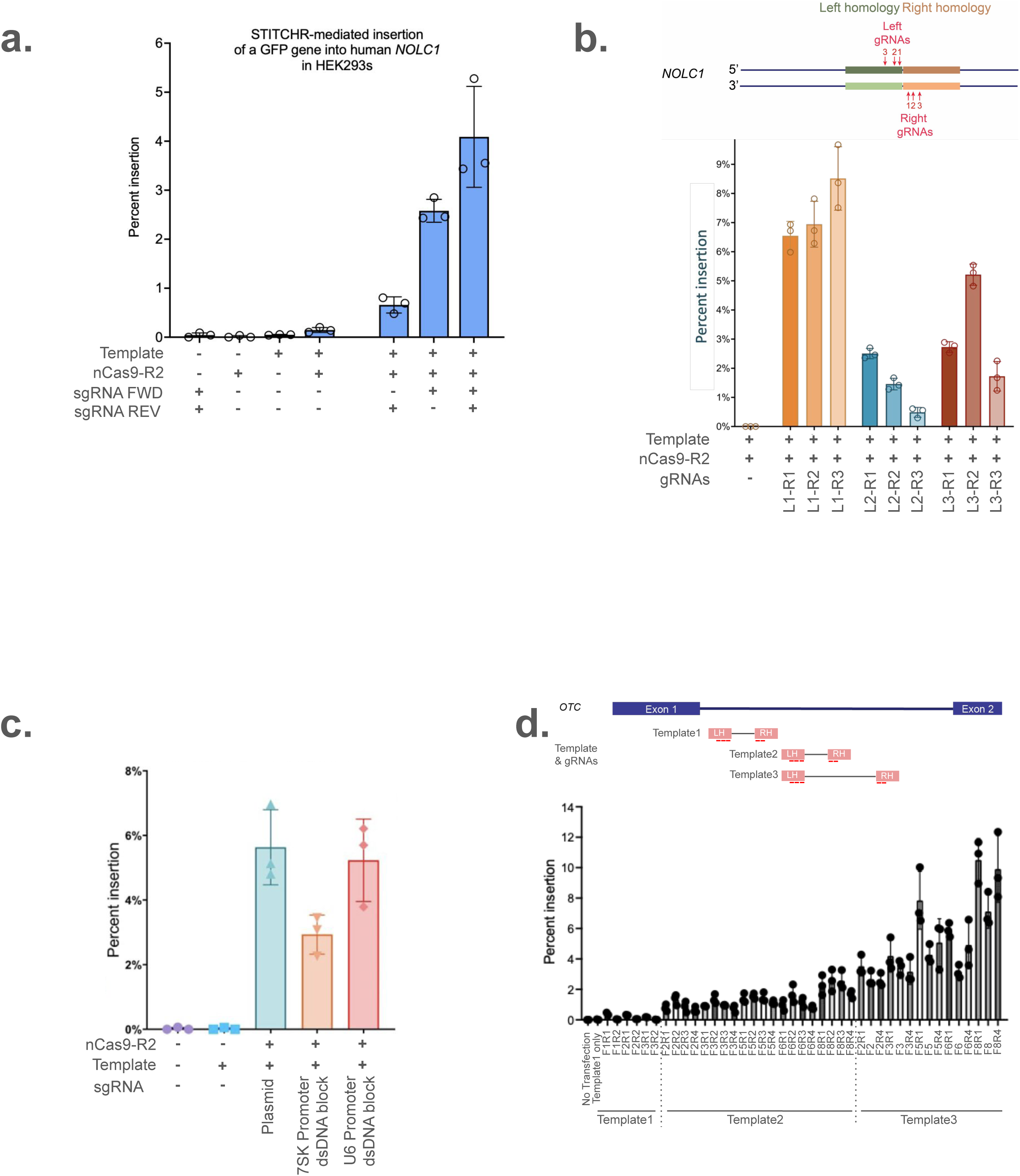
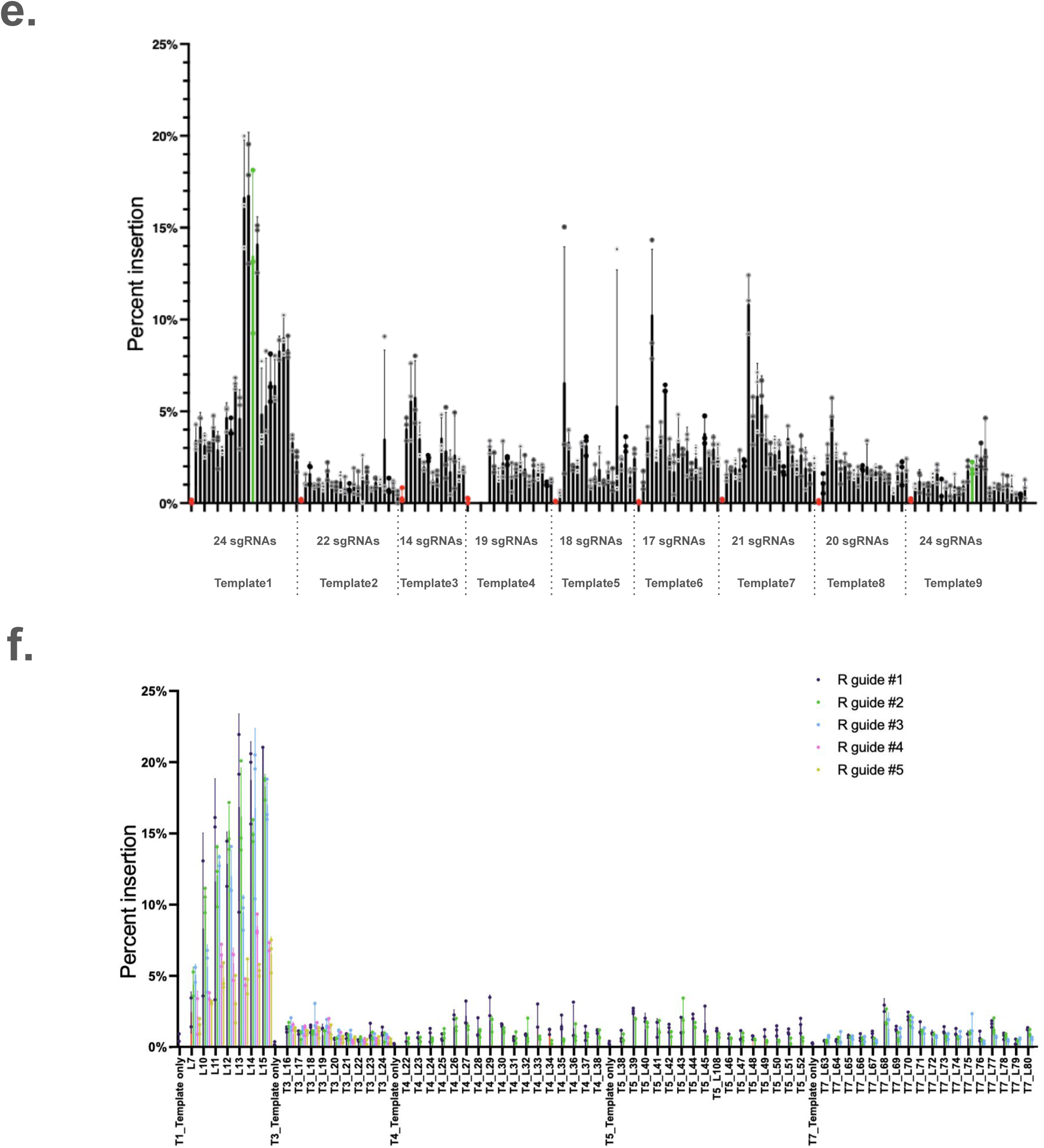
Systematic sgRNA screening identifies optimal guide combinations for efficient T-PGI-mediated insertion. (a) ddPCR analysis of GFP insertion at the *NOLC1* locus using forward (left), reverse (right), or paired sgRNAs. (b) Screening of sgRNA pairs targeting the left and right homology arms at *NOLC1 locus*. Top: schematic of guide positions. Bottom: insertion efficiencies for all pairwise combinations measured by ddPCR. (c) Comparison of sgRNA expression strategies. Linear dsDNA templates driven by U6 or 7SK promoters were tested against plasmid-based expression, and insertion efficiencies were quantified by ddPCR. (d) Combinatorial guide-template screen at the *OTC* locus. Top: three donor templates targeting intron 1 of *OTC* gene. Bottom: ddPCR-based efficiencies for 41 guide–template combinations. (e) First-round screen of *CAR19* insertion at the *AAVS1* locus. Nine donor templates were paired with a single sgRNA; Red dots indicate template-only controls; green box highlights Template 1/9 with sgRNA L15. (f) Second-round screen evaluating dual-sgRNA combinations across the top five donor templates for *CAR19* insertion at *AAVS1* by ddPCR measurement. All experiments were performed in HEK293 cells.

To further refine guide selection, we performed a combinatorial screen at the *NOLC1* locus using three sgRNAs per homology arm. Pairwise combinations showed notable variability in editing efficiency, underscoring the need for empirical sgRNA optimization (**Fig. 4b**). To streamline large-scale guide screening, we developed an approach that would eliminate the need for time-consuming cloning or costly synthetic sgRNAs. To this end, we tested whether sgRNA cassettes could be expressed from short, linear double-stranded DNA (dsDNA) fragments encoding either a *U6* or *7SK* promoter upstream of the guide sequence. The feasibility of this strategy was motivated by the compact nature of RNA polymerase III–driven promoters such as *U6* and *7SK*, which support efficient transcription initiation and defined termination within short sequences, ideal for dsDNA delivery. The *U6*-driven dsDNA format yielded insertion efficiencies comparable to conventional plasmid-based sgRNA expression, while the *7SK*-driven format achieved ∼50% of that activity (**Fig. 4c**). This system enabled rapid combinatorial testing of sgRNA–template pairs in downstream applications.

We applied this approach to integrate full-length *OTC* cDNA at its endogenous locus, combining three donor templates with various sgRNA pairs, yielding 41 total combinations. Insertion efficiencies ranged from <0.1% to >10%, confirming the importance of guide–template pairing (**Fig. 4d**). Finally, to demonstrate the utility of this screening strategy in a clinically relevant context, we performed a two-stage screen to insert a *CAR19* transgene at the *AAVS1* locus. To identify the optimal combination of template, homology arm design, and sgRNA configuration, we first designed 9 candidate donor templates and tested them in conjunction with 155 single-guide RNAs targeting flanking regions. Based on this initial screen, we selected the top five performing templates (Template1, 3, 4, 5 and 7) for further analysis (**Fig. 4e**). In the second stage, we systematically tested 200 dual-guide combinations across these five templates. This comprehensive approach enabled the identification of an optimal configuration, template 1 combined with the sgRNA pair L15 and R7, which yielded the highest insertion efficiency (**Fig. 4f**). Intriguingly, we also found that the direction of the TRPT reaction markedly influenced editing outcomes: template 1 and template 9 contained identical sequences but were written in opposite orientations. Despite using the same L15 sgRNA, template 1 achieved ∼13% insertion efficiency, whereas template 9 reached only ∼3%, underscoring that writing direction is a critical determinant of genome editing efficiency (**Fig. 4e**). These results collectively underscore the importance of template design, particularly the positioning of homology arms, as well as the precise location, pairing of sgRNAs, and reaction directionality. The ability to fine-tune these parameters was critical for maximizing editing outcomes at the *AAVS1* locus and serves as a generalizable principle for applying T-PGI to other genomic targets.

### Protein engineering improves core enzyme performance

In Fell et al. (Fell et al., 2025), the nickase Cas9 and R2 retrotransposon proteins were expressed as a single polypeptide through direct fusion, providing an initial proof of concept. However, whether this configuration represented the most effective architecture for enzymatic activity remained unclear. To systematically optimize the nCas9–R2 core module, we evaluated several engineering strategies. First, we verified that fusion orientation was critical: while the original nCas9–R2 configuration supported insertion activity, reversing the order (R2–nCas9) markedly reduced efficiency (**Fig. 5a**). We next examined 17 linker variants spanning flexible linkers, rigid linkers, and self-cleaving peptides. Among these, bicistronic expression mediated by P2A outperformed most direct fusions, yielding approximately two-fold higher insertion efficiencies (**Fig. 5b**). Given the importance of nuclear localization for Cas9 function, we tested whether additional nuclear localization signals (NLSs) could enhance activity. Introducing NLSs at both termini of R2 increased insertion efficiency by 4–5-fold, whereas removal of both NLSs abolished activity (**Fig. 5c**). Finally, we compared bicistronic expression (via T2A or P2A) with a separate protein expression system. All strategies achieved comparable activity, demonstrating that enforced fusion can compromise performance, while independent expression maintains robust activity (**Fig. 5d**)

**Fig 5.**
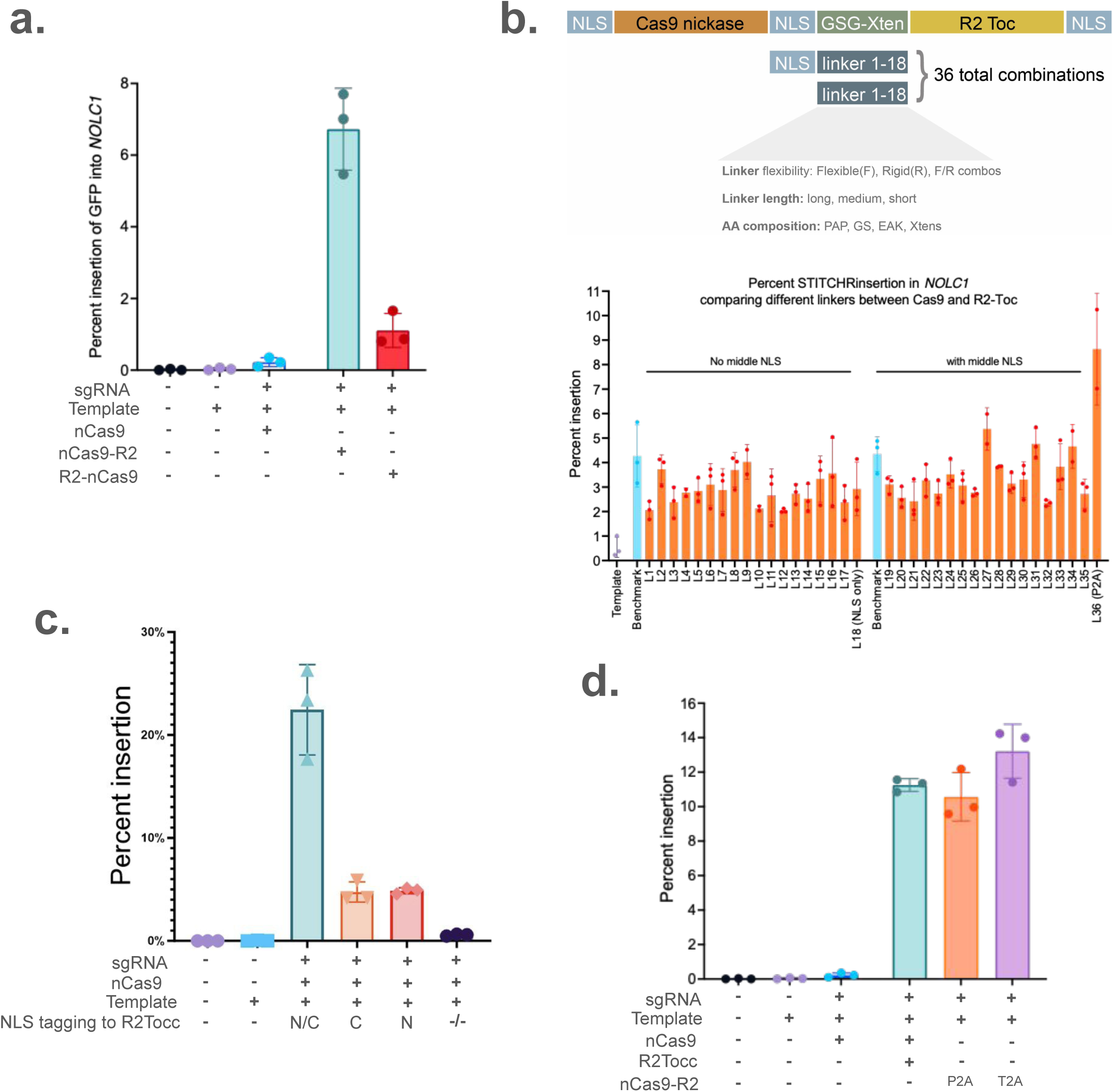
Protein engineering improves enzyme configuration, linker design, and nuclear localization for enhanced T-PGI activity. (a) Comparison of fusion orientations. The nCas9–R2 configuration enables robust insertion, while the reverse (R2–nCas9) markedly reduces efficiency. (b) Screening of 36 linker variants between nCas9 and R2, with or without an upstream nuclear localization signal (NLS). (c) Assessment of NLS tagging on R2 activity, showing enhanced insertion with dual-terminal NLSs. (d) Comparison of expression strategies: separate plasmid co-expression, T2A linkage, and P2A linkage. All experiments involved *GFP* insertion at the *NOLC1* locus in HEK293 cells, with insertion efficiency quantified by ddPCR.

### Terminal and internal modifications influence enzyme stability and activity

Given the low steady-state levels of R2Tocc compared to nCas9, we sought to improve protein expression and stability through terminal modifications (**Supp Fig 3a**). We first tested the addition of STABILON (Rethi-NAgy et al., 2022) and HiBiT tags at the C-terminus of R2Tocc, which increased detectable protein levels by approximately 50% (**Supp Fig 3a**). However, despite the increased expression, insertion efficiency was reduced by more than half, indicating that the C-terminus of R2 plays a critical functional role and is likely sensitive to steric hindrance or structural interference (**Supp Fig 3b**). Supporting this notion, we observed that nCas9 protein levels were approximately 20-fold higher than R2Tocc under identical expression conditions, suggesting that R2 is subject to rapid turnover or degradation (**Supp Fig 3a**).

We also evaluated N-terminal tagging by fusing a VP64 transactivation domain or the IDR domain of FUS to the R2 enzyme. Similar to the C-terminal modification, this N-terminal fusion also resulted in a dramatic reduction in insertion efficiency, further supporting the hypothesis that both termini are functionally constrained (data not shown). Taken together, these results suggest that conventional approaches for enhancing protein expression or stability, such as terminal epitope tagging, may not be applicable to R2 enzymes, and that alternative strategies, including internal engineering, are needed to enhance activity without disrupting function.

### Internal protein engineering reveals flexible regions and enables functional enhancement

To circumvent the limitations of terminal tagging, we turned to internal engineering of the R2 enzyme to identify flexible regions that could be modified without compromising enzymatic function. We focused on a ∼250 amino acid intrinsically disordered region (IDR) between the ZNF and RT domains. Based on structural predictions, this region was hypothesized to lack stable folding and therefore might tolerate deletions or insertions. We generated a series of truncation mutants and discovered that deleting a specific 50 amino acid segment between the ZNF domain and a putative RNA-binding region (termed Deletion 4) not only preserved activity but enhanced insertion efficiency (**Fig. 6a**). This suggests that the deleted region is dispensable for reverse transcription and may have previously hindered proper folding or activity.

**Fig 6.**
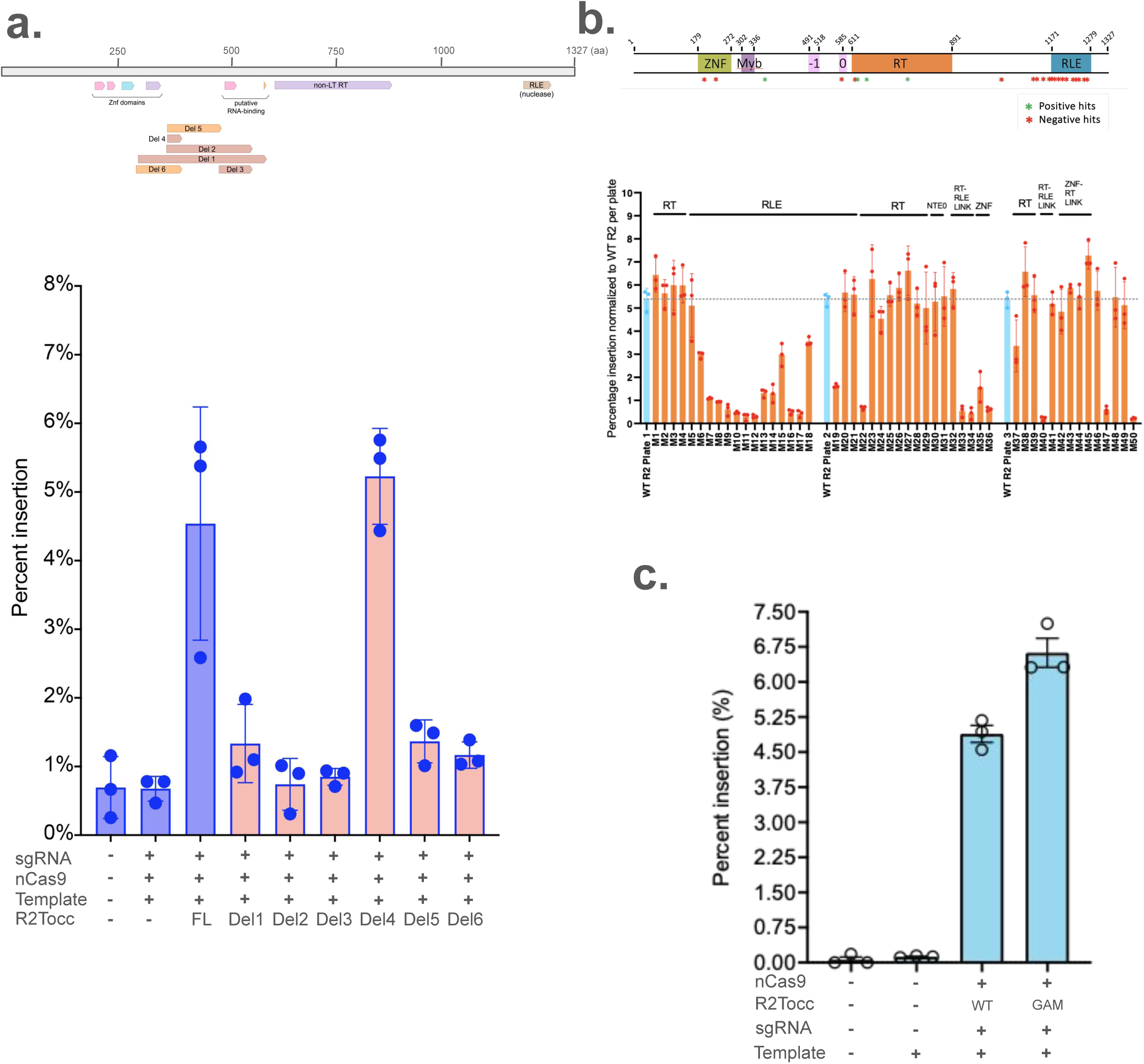
Internal protein engineering of R2Tocc affects activity and enables functional modularization. (a) Internal deletion screen across a ∼250 amino acid intrinsically disordered region (IDR) between the ZNF domain and RT domains. Top: schematic of deletion mutant locations. Bottom: ddPCR-based insertion efficiency of each R2Tocc variant. (b) Functional assessment of point mutations in conserved domains of R2, including RT, RLE, ZNF, and the RT–RLE linker, or insertion mutants into Del4 regions. Top: mapping of mutations with increased or decreased insertion efficiency. Bottom: ddPCR analysis of each mutant. (c) Insertion of the Mu Gam protein into the Deletion 4 site resulted in higher insertion efficiency, highlighting the potential for internal modularization. All experiments were performed at the *NOLC1* locus using GFP insertion templates, with efficiency quantified by ddPCR.

In parallel, we introduced point mutations into highly conserved residues or RNA contacting residues within known functional domains of the R2 enzyme, including the reverse transcriptase (RT), RNase-like endonuclease (RLE), and zinc finger (ZNF) regions, as well as the RT-RLE linker. These mutational scans helped delineate regions that are critical versus those that are tolerant to perturbation, informing future design constraints (**Fig. 6b**).

Finally, having identified Deletion 4 as a permissive site, we tested whether this region could serve as an internal docking site for protein domain insertions to further improve activity or stability. We engineered three constructs by inserting: (1) XTEN Linker to the unstructured region (M43), (2) an ssDNA-binding domain to potentially stabilize TPRT intermediates (M44), and (3) the Mu Gam protein from bacteriophage, known to bind and protect free DNA ends (Di Fagagna et al., 2003). While the XTEN linker and ssDNA-binding domain yielded modest or no improvement, insertion of Mu Gam led to a ∼30% increase in insertion efficiency (M45 in the mutant screening) (**Fig. 6b and c**). These results demonstrate that internal regions of R2 can tolerate insertions and be repurposed to introduce functional modules, providing a strategy for enhancing activity through domain engineering.

## Discussion

In this study, we present a platform-level engineering effort that transforms the initial version of STITCHR into a more efficient, accurate, and modular genome editing tool; we refer to this engineered implementation as T-PGI (Transposon-based Programmable Genome Insertion). Like the original STITCHR concept, T-PGI enables RNA-templated, scarless insertion of DNA cargo via target-primed reverse transcription (TPRT) without relying on double-strand breaks. By systematically optimizing each core component of the system, including the retrotransposon enzyme, donor template design, guide RNA strategy, and expression architecture, we addressed key limitations of the original platform and expanded its potential utility for therapeutic genome engineering. Throughout this manuscript, “STITCHR” denotes the original implementation, whereas “T-PGI” denotes the engineered system described here.

A major motivation for this work stemmed from the observation that the original STITCHR enzyme, R2Tg, exhibited unwanted integration into its native target site, the 28S rDNA locus. Even without perfect homology, this off-target activity posed a risk for therapeutic application. Our decision to adopt R2Tocc as the core enzyme was based on prior studies showing its markedly reduced background activity at 28S rDNA, and we confirmed that it performs comparably to R2Tg in terms of insertion efficiency (Fell et al., 2025). This shift reflects a general principle in genome editing, namely, that safety and specificity are as critical as raw efficiency when moving toward clinical translation.

We next addressed the challenge of cargo flexibility. Given that R2-mediated reverse transcription is prone to 5’ truncations, particularly for long templates, it was unclear whether T-PGI could accommodate therapeutically relevant cargo sizes. Through systematic testing, we demonstrated successful integration of inserts ranging from 26 bp to over 4 kb without major loss in efficiency. Furthermore, we showed that T-PGI can support large-scale sequence replacement, achieving precise cassette insertion following deletions up to 5 kb in size. These findings confirm that TPRT-mediated genome editing can be applied not only to small corrections but also to full gene replacement or exon-level repair.

Insertion fidelity is another critical aspect for translational genome editing platforms. We conducted both short- and long-read sequencing to assess the accuracy of T-PGI-mediated integration. While editing precision was high overall, we did observe a modest decrease in sequence fidelity as cargo length increased, a trend consistent with other reverse transcriptase-based systems (Anzalone et al., 2021). Importantly, we detected no increase in local indels, scars, or concatemer formation, suggesting that even with partial sequence loss, the integration process remains clean and localized. These results highlight the value of pairing TPRT with rational template design and nickase-based targeting to maximize fidelity.

Our study also emphasizes the importance of donor template configuration and sgRNA positioning. Template elements such as UTRs, homology arm length, and post-insertion PAM removal were all found to influence efficiency. Through large-scale combinatorial template and sgRNA screening, including a clinically relevant test case at the *AAVS1* locus, we showed that dual-guide designs and precise arm placement are key determinants of editing success. The ability to express sgRNAs from linear dsDNA cassettes further enhances the platform’s flexibility and makes high-throughput screening more tractable. We also note that insertion efficiencies were occasionally variable across replicates, largely reflecting differences in transfection efficiency and cellular status in HEK293 cells. This source of variability underscores the importance of cellular context and delivery optimization when evaluating transfection-based genome editing.

Beyond guide and template design, we demonstrate that enzyme configuration plays a major role in editing outcomes. While the original STITCHR used a direct nCas9–R2 fusion, our comparison of different linker types, expression systems, and nuclear localization signals revealed that a P2A-based co-expression strategy and dual-terminal NLS tagging significantly improve performance. These improvements likely stem from enhanced nuclear availability and reduced steric hindrance.

Finally, internal engineering of the R2 enzyme proved to be a powerful strategy for improving activity. By targeting an intrinsically disordered region between the ZNF and RT domains, we identified a deletion (Deletion 4) that enhances insertion efficiency without disrupting core function. We further showed that this region can tolerate insertion of modular elements, such as the Mu Gam protein, which conferred a ∼30% improvement in editing efficiency, likely by protecting the dsDNA-end during the genome editing. These results position R2 as a structurally flexible and engineerable scaffold for future protein enhancement.

Together, our findings establish a revised version of STITCHR that is substantially more efficient, accurate, and tunable than its predecessor. The ability to combine programmable targeting, cargo flexibility, and modular enzyme design opens new opportunities for therapeutic genome editing across a wide range of disease models and cell types.

## Materials and Methods

### Cell culture and transfection

HEK293T cells (ATCC) were maintained in DMEM (11965092, Gibco) supplemented with 10% FBS (A3160501, Gibco). Cells were dissociated using 0.25% Trypsin-EDTA (15400054, Gibco) and seeded into 96-well tissue culture plates at a density of 30,000 cells per well. After 16 hours, the medium was replaced with Opti-MEM (31985070, Thermo Fisher Scientific), and cells were transfected at approximately 70% confluency. For each well, the transfection mix contained 0.4 μL of Lipofectamine 2000 (11668019, Thermo Fisher Scientific) and a defined amount of plasmid DNA. Sixteen hours post-transfection, the medium was switched back to DMEM containing 10% FBS, and cells were cultured for an additional three days. For sgRNA expression from dsDNA oligos (See **TableS1**), 1 to 10 ng of dsDNA was transfected per well in a 96-well format. You can find the homology arm, mutant, linker and template sequences in **TableS1**.

### Double-stranded oligo-based sgRNA expression system

To express the sgRNA, a 408 bp double-stranded DNA fragment was synthesized (IDT). The construct includes a *U6* promoter (250 bp) or *7SK* promoter (268 bp), a 20 bp spacer sequence, and an 83 bp gRNA scaffold and termination sequence.

### sgRNA design

sgRNA sequences were designed using the CRISPR design module in Benchling (Benchling, San Francisco, CA), selecting guides with high predicted on-target efficacy and low off-target scores.

### gDNA isolation

Cells were lysed using QuickExtract DNA Extraction Solution (Lucigen, QE0905T), and genomic DNA was subsequently purified by SPRI-based magnetic bead cleanup.

### Digital droplet PCR (ddPCR)

Custom primers and probes were designed to measure editing in all referenced loci. Results were normalized to custom reference assays targeting unedited regions of the same genes in the respective species. Probes were dual labelled with 3′-3IABkFQ and either 5′-carboxyfluorescein (FAM) for edit targets or 5′-hexachloro-fluorescein phosphoramidite (HEX) for reference. Assays were validated using gBlocks representing edit outcomes to test for both specificity and linearity. All primers, probes, and gBlocks were synthesized by IDT. Each reaction contained 12 µL of 2x ddPCRSupermix for probes (No dUTP) (1863025 Bio-Rad), 1.2 µL of each primer and probe mix to final concentration of 0.5 uM for each primer and 0.25 uM for each probe, 0.12 µL each of HindIII and Eco91I (FD0505 and FD0394, Thermo Fisher Scientific), 10-20 ng of DNA and water to a final volume of 24 µL. Droplets were generated on the AutoDG Instrument for automated droplet generation (186410, Bio-Rad). PCR amplification was performed with the following cycling parameters: initial denaturation at 95 °C for 10 min, followed by 40 cycles of denaturation at 94 °C for 30 s and combined annealing/extension step at 58 °C for 1 min, and a final step at 98 °C for 10 min. Data acquisition and analysis were performed on the QX200 Droplet Reader.

### Hibit assay

HiBiT expression was measured using the Nano-Glo HiBiT Lytic Detection System (N3050, Promega, Madison, WI, USA) according to the manufacturer’s instructions. A 50 μL master mix, prepared by combining Nano-Glo HiBiT Lytic Buffer, Substrate, and LgBiT Protein at a 100:2:1 ratio, was added to each cell well. Plates were shaken for 10 minutes and then incubated for an additional 10 minutes before luminescence was measured on a GloMax Explorer (GM3500, Promega) with a 0.3-second integration time. For in vivo studies, 125 μL of master mix was added to 25 μL of serum, with all other steps performed identically.

### Cell-Titer Glow assay

To quantify the cell viability, CellTiter-Glo® 2.0 Luminescent Cell Viability Assay (Promega) was used. Cells were plated with at least two rows as a gap to avoid signal cross-contamination. For each well in a 96-well black wall plate, approximately 75 μL of CellTiter-Glo® 2.0 was added to the media. The plate was shaken at 600 rpm at room temperature without light exposure for 10 min before reading plate luminescence using a Paradigm plate reader.

### Amp-seq and Long-read sequencing

Target regions were amplified using Q5 Hot Start High-Fidelity 2× Master Mix (NEB, M0494X) for 25 cycles. Annealing temperatures were determined based on the gene-specific portion of the primers using NEB’s online Tm calculator (https://tmcalculator.neb.com/). PCR products were either analyzed by electrophoresis on a 2% agarose gel or used directly as input for a second-round Illumina barcoding PCR. For Oxford Nanopore sequencing, libraries were prepared according to the manufacturer’s instructions using the Native Barcoding Kit 96 V14 (Oxford Nanopore, SQK-NBD114.96). PacBio HiFi sequencing was performed by Novogene as a commercial service.

### Data analysis

Raw signals from Oxford Nanopore Technology MinION were basecalled and demultiplexed using Dorado v0.7 (https://github.com/nanoporetech/dorado) using the *sup* model. Porechop (https://github.com/rrwick/Porechop) was used to trim the adapters, and chopper (https://github.com/wdecoster/chopper) was used for quality control of the reads by removing reads shorter than 300 bp and average quality of less than 10. Nanoplot (https://github.com/wdecoster/NanoPlot) was used to generate read quality plots. Filtered reads were then mapped to the reference sequence using bwa mem with parameters -x ont2d. Alignment files were visualized using Integrative Genomics Viewer (IGV), and alignment metrics were calculated using samtools v1.18.

PacBio HiFi reads were mapped to the reference sequence using minimap2 with parameters -ax map-hifi. Alignment files were visualized using IGV, and fidelity statistics were calculated using custom scripts by extracting insertions, deletions, and mutation information.

Illumina sequences were analyzed using CRISPResso2, and fidelity statistics were reported based on the insertions, deletions, and/or mutation information.

---

**Supplementary Fig 1.**
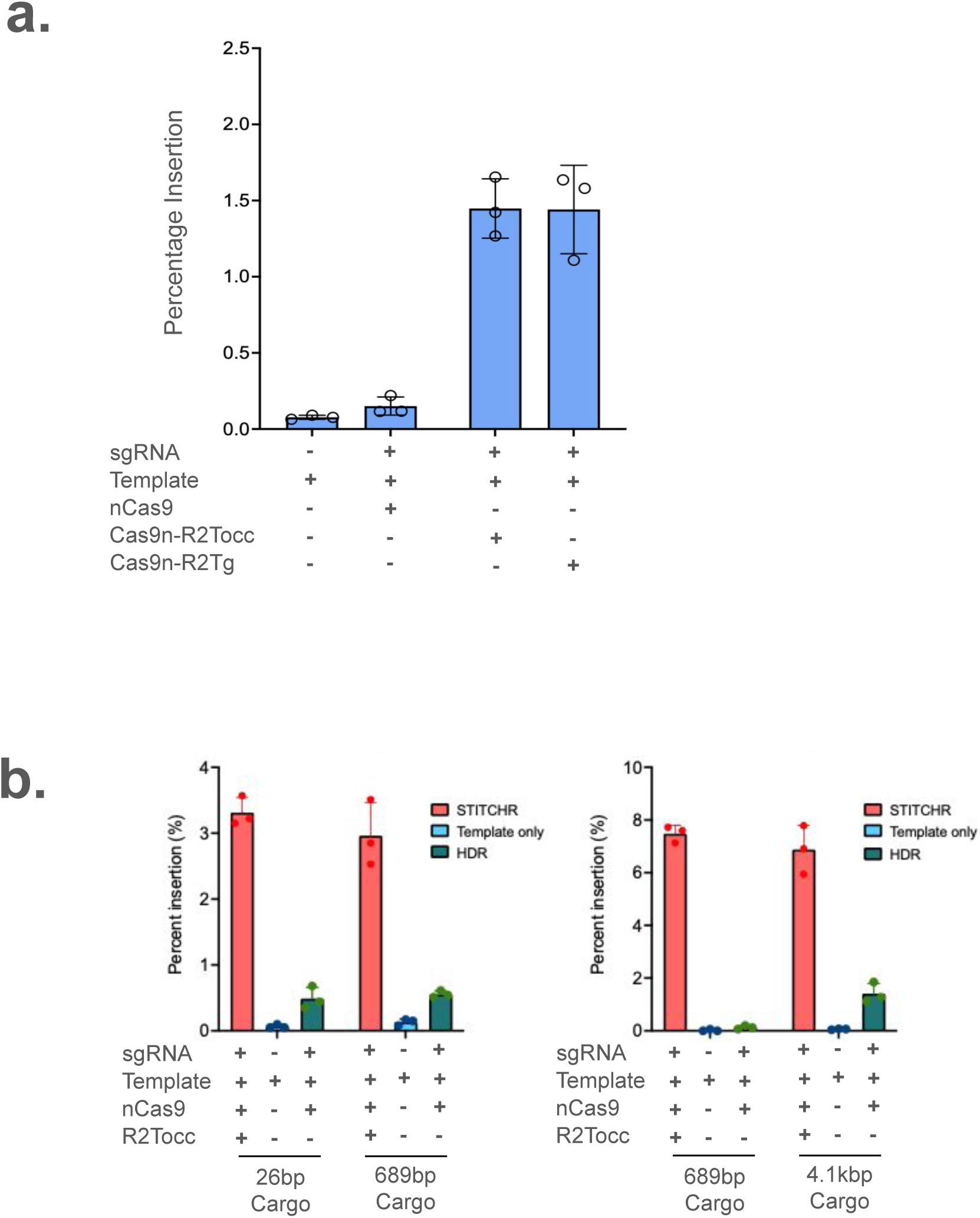
Comparison of R2Tg and R2Tocc and insertion efficiency across cargo sizes. (a) ddPCR-based quantification of *Gluc* gene insertion at the *AAVS1* locus using R2Tg or R2Tocc with a shared donor in HEK293 cell. Negative controls include template only or a combination of Cas9 nickase (nCas9), sgRNAs, and donor template (HDR-like condition). (b) ddPCR analysis of R2Tocc-mediated insertion at the *NOLC1* locus using donor templates of 26 bp, 689 bp, or 4.1 kb. Controls included template only and HDR-like condition.

**Supplementary Fig 2.**
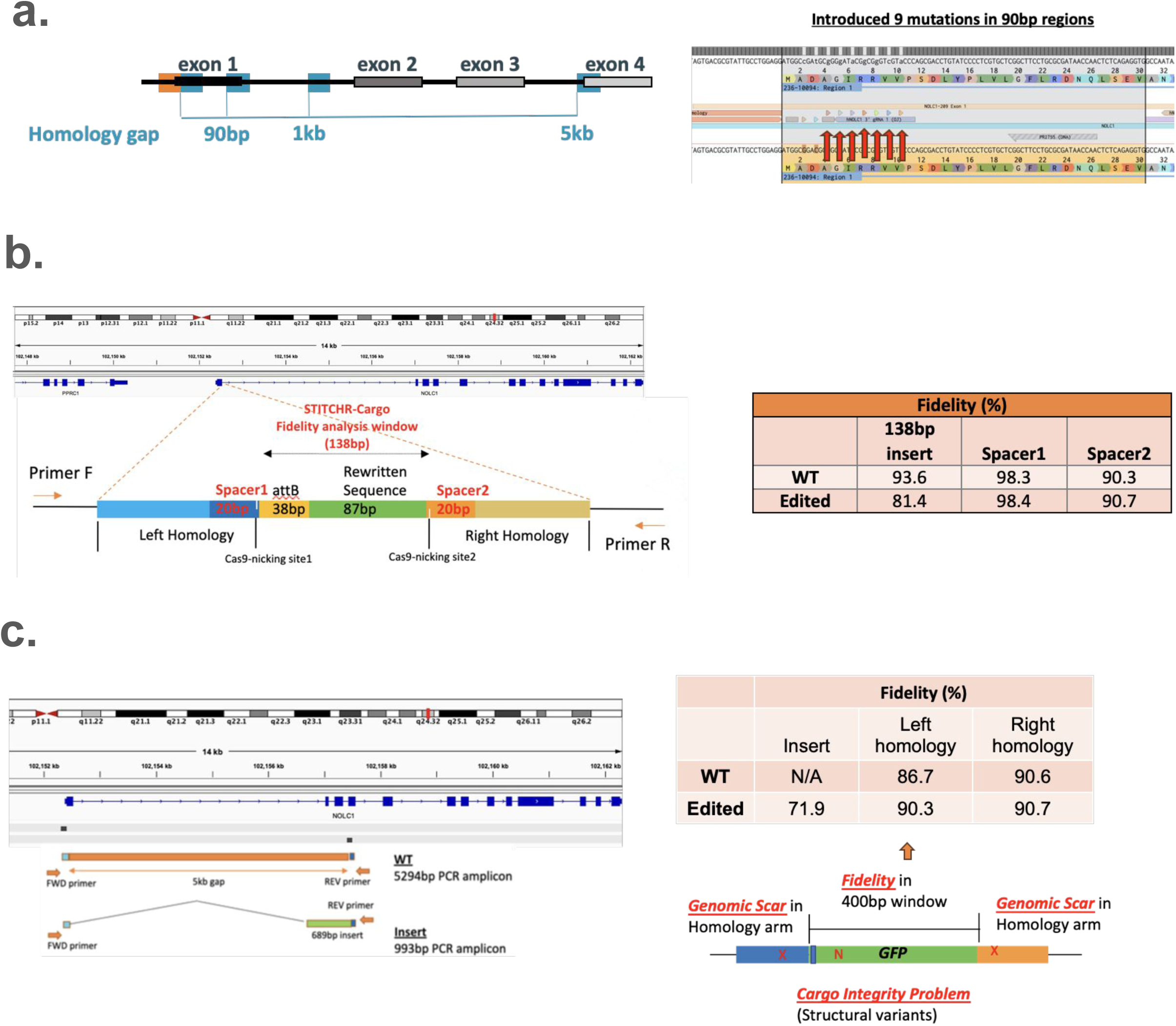
NGS-based assessment of T-PGI precision and integrity. (a) Left: Schematic of donor design and homology arm placement for precision rewriting of a 90 bp genomic region. Right: Targeted amplicon sequencing confirms accurate incorporation of 9 programmed missense mutations. (b) Left: Schematic of the engineered locus showing insertion of a 38 bp attB and 87 bp rewritten sequence.Right: Amplicon-based Illumina sequencing analysis of editing fidelity across the 138 bp insertion and flanking regions, including Cas9 nicking sites and two 20 bp spacer sequences (Spacer 1 and 2). (c) Left: Genome browser view showing edited allele with 5 kb deletion replaced by 689 bp GFP. Right: PacBio HiFi sequencing-based fidelity assessment across a 400 bp window within the inserted GFP cassette (excluding homopolymers) and both homology arms. No structural variants or aberrant insertions were detected.

**Supplementary Fig 3.**
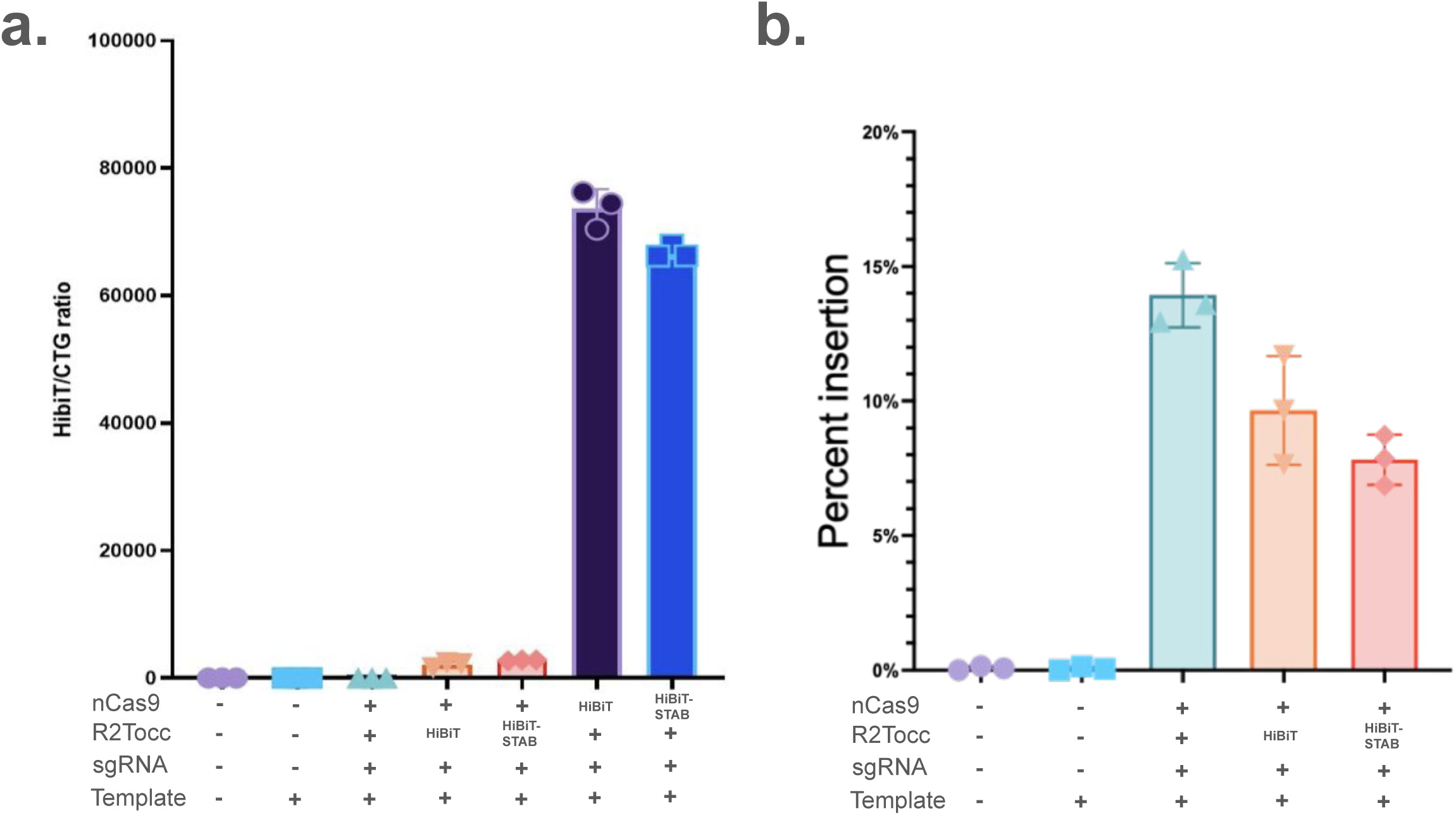
Tagging strategies impact R2Tocc stability and insertion efficiency. (a) Expression analysis of R2Tocc with or without C-terminal STABILON and HiBiT tags, measured by HiBiT activity normalized to CellTiter-Glo (CTG) luminescence. STABILON tag (STAB) increased R2Tocc expression by ∼50%, but had a minor effect on nCas9. R2Tocc protein levels remained ∼20-fold lower than nCas9. (b) Functional assessment of tagged R2Tocc variants. Despite higher expression, C-terminal tagging reduced GFP insertion efficiency by over 50%, as quantified by ddPCR at the *NOLC1* locus.

